# Visual mapping on the brain - caveats for multifocal fMRI

**DOI:** 10.1101/676023

**Authors:** David P. Crewther, Shaun A. S. Seixas, Sheila G. Crewther

## Abstract

While multifocal electroretinography has become a standard ophthalmological technique, its use in cortical neuroimaging has been lesser. Vanni et al. (2005) presented the first exploration of the multifocal visual mapping methodology with fMRI. This commentary confirms the utility of this method, but also presents empirical results which suggest caveats for the use of the technique. In the current study rapid multifocal fMRI was established using m-sequence pseudo-random binary stimuli applied to visual field mapping in six young adults with normal vision. Nine contiguous regions of visual field – two rings of 4 patches with a central patch, areas scaled for cortical magnification, were pseudo-randomly stimulated, with patterned or grey images. The decorrelation of stimulus patches allowed all 256 volumes to be used for the analysis of each of the nine stimulus areas. Strong localized activation was observed for each of the four peripheral regions with the location of the activation conforming to the expected visual field retinotopy. The inner regions, including the foveal patch, did not significantly activate. We propose, on the basis of a simple correlational model of simulated eye movements, that the loss of signal is due to gaze instability. Thus, while the rapid multifocal method can be successfully applied to fMRI, the results appear quite sensitive to eye movements, the effects of which may have been overlooked by smoothing evoked responses to achieve a retinotopic map.

## Introduction

The standard fMRI procedure for retinal mapping involves stimulation with a slowly rotating flickering wedge or expanding annulus (Sereno et al., 1995;DeYoe et al., 1996;Engel et al., 1997). The position in the visual field is then extracted as the phase of the Fourier transformed time series for each voxel in visual cortex. This allows the boundaries between cortical areas to be accurately located (Engel et al., 1997), but cannot eliminate positional errors in the phase encoding process contributed by the spatial and temporal aspects of the BOLD signal, as well as noise. Vanni et al. (2005) suggested that an alternate approach, stimulating multiple discrete regions of visual field and using the peak activation centres to form a grid map, was preferable for efficient mapping.

While visual field regions can be sequentially mapped with individual focal stimuli (Schneider et al., 1993) the resultant map requires inordinate scanning time. Although event-related functional magnetic resonance imaging (erfMRI) allows for randomised stimulus sequences and investigation of independent hemodynamic responses (HDR), it traditionally lacks the statistical power of the block design (Liu et al., 2001). By comparison, a combination of event-related and block design fMRI results in an erfMRI paradigm (m-sequencing) that allows for the extraction of stimulus responses while retaining acceptable statistical power (Liu, 2004).

Historically, multifocal m-sequence paradigms were developed as a tool for clinical electrophysiology to allow more efficient mapping of visual field topological pathologies (Hood, 2000). Mapping is achieved by sectoring the visual field into discrete (multifocal) elements or patches whose temporal stimulation sequences are independent and decorrelated (provided the neuronal memory is sufficiently small) (Sutter, 1992). Therefore, in a multifocal m-sequence protocol every iteration of the stimulus sequence adds an equal amount of statistical power to each of the stimulus elements, giving maximal power and efficiency (Sutter, 1992).

The most common white-noise implementations use m-sequence binary stimuli (suitable for the comparison of two states, e.g., ‘on’ and ‘off’) with pseudo-random sequences generated from primitive polynomials, modulo 2 (Press et al., 1986), that allow analysis through convolution with the regenerated sequence. The efficiency of such sequencing techniques is remarkable. Four minutes of recording is sufficient to produce reliable electroretinographic (ERG) or visual evoked potential (VEP) waveforms from over 60 separate locations in visual field (Sutter, 1992). The capacity to access the non-linear structure of the visual response is also an advantage of the m-sequence technique over other similar decorrelational methods (Maddess et al., 1997). For example, the second order kernels of the flash VEP have been related to the contributions of the magnocellular and parvocellular pathways (Klistorner et al., 1997;Crewther et al., 1999).

Although the m-sequence has proven itself over the past decade to be a very useful tool in electrophysiology, its transition to fMRI has occurred rather slowly. The neuroimaging literature contains only a few comments on the statistical attributes of the m-sequence (Liu et al., 2001;Buracas and Boynton, 2002;Liu, 2004). Subsequently, a small number of experimental utilizations of m-sequence methods have appeared (Kellman et al., 2003;Hansen et al., 2004;Vanni et al., 2004;de Zwart et al., 2005;Henriksson et al., 2012).

Kellman et al. (2003) investigated the temporal non-linearity of BOLD responses to patterned visual stimuli with temporal modulation obeying a pseudo-random m-sequence. Their findings demonstrated a diminution of second order BOLD responses in V1 with increasing interstimulus intervals. Differences were also seen between foveal and peripheral nonlinearities. Hansen et al. (Hansen et al., 2004) used independent m-sequence derived spatial stimuli to investigate the spatial linearity (additivity) of V1 BOLD responses in two participants and found that positive BOLD responses in V1 were adequately characterized by linear spatial summation. While both of these studies utilized the power of m-sequence protocols for independent responses, neither investigation fully captured the multifocal capability of m-sequence stimuli for spatial mapping. M-sequence paradigms have also been used to investigate such problems as the temporal limitations of the BOLD response (de Zwart et al., 2005), fMRI seeded electroencephalogram (EEG) source localization (Vanni et al., 2004) and visual evoked magnetic field source localization (Tabuchi et al., 2002;Nishiyama et al., 2004).

Vanni et al. (2005) used a multiple linear regression multifocal technique - the pattern-pulse method (James, 2003), for retinotopic mapping with fMRI. They reported the capacity to simultaneously map 60 independent regions of visual field. As with the visualisation of phase encoded mapping methods, Vanni et al. (2005) presented their findings on smoothed grey matter segmented cortical surfaces. A grid of 30 points was positioned on each hemisphere with vertices corresponding to the peak activation from each of the patches. These visualisations show a predominant mapping of visual cortex around the calcarine sulcus (V1 & V2) and appear comparable to phase encoded maps. The authors reported that the projection of voxel activations to the segmented cortical surface produced patched responses which required voxel clustering thresholds and spatial smoothing for contiguous mapping. They ascribed the patchiness to both uneven assignment of voxels and spatial distortions from the cortical unfolding process.

Baker et al. (2006) proposed a multimodal EEG/fMRI approach. In this study the investigators co-localized the BOLD response with EEG (VEP) dipole sources in two participants. Interestingly the fMRI stimulation involved the standard phase encoded method (rotating wedge, expanding annuli), whereas EEG stimulation was via a multifocal m-sequence (60 stimulus regions), acquired in a separate session. The continuous fMRI field maps were interpolated to fit the VEP stimulus regions. Results showed an approximate level of BOLD/VEP co-localized source response. Henriksson et al. (2012) compared the standard topography methods with multifocal methods and demonstrated similar capability.

In the light of the existing literature, the current investigation applied the standard m-sequence multifocal method to fMRI aiming to register functional integrity across the visual field in addition to retinotopic mapping. A very rapid event related protocol - nearly three times as rapid as that of Vanni et al. (2005), was employed. Sensitivity of such a protocol to simulated eye movements suggests limitations in the application of multifocal methods to fMRI scanning as currently implemented.

## Materials and Methods

### Subjects

Data was acquired from six participants (1 male, 5 female; mean age = 25 yr) who gave voluntary consent, with the research program approved by the relevant institutional ethics committee. Exclusion criteria included any history medical or psychiatric disorders, neurosurgery or metal implants. Participants were asked to view the stimulus sequences while maintaining fixation on the centre of the stimulus, rear projected on a screen mounted at the participant’s feet and observed via a mirror mounted within the head coil helmet.

### Stimulus Presentation

The stimulus consisted of a circle divided into 9 regions: a central foveal region, four central quadrants and four peripheral quadrants (see Figure 1a). The total stimulus subtended 8 degrees of visual field. The central foveal region was 0.5 deg across. The patches of the first ring of quadrants were centred at 0.6 deg eccentricity. The peripheral patches were approximately 6 times larger (in visual field), centred at 2.6 deg eccentricity. Using the magnification factor function derived from Duncan and Boynton (Duncan and Boynton, 2003), the patches in the middle and peripheral rings stimulated approximately equal areas (∼ 225 mm^2^) on the cortical surface of area V1.

**Fig. 1.**
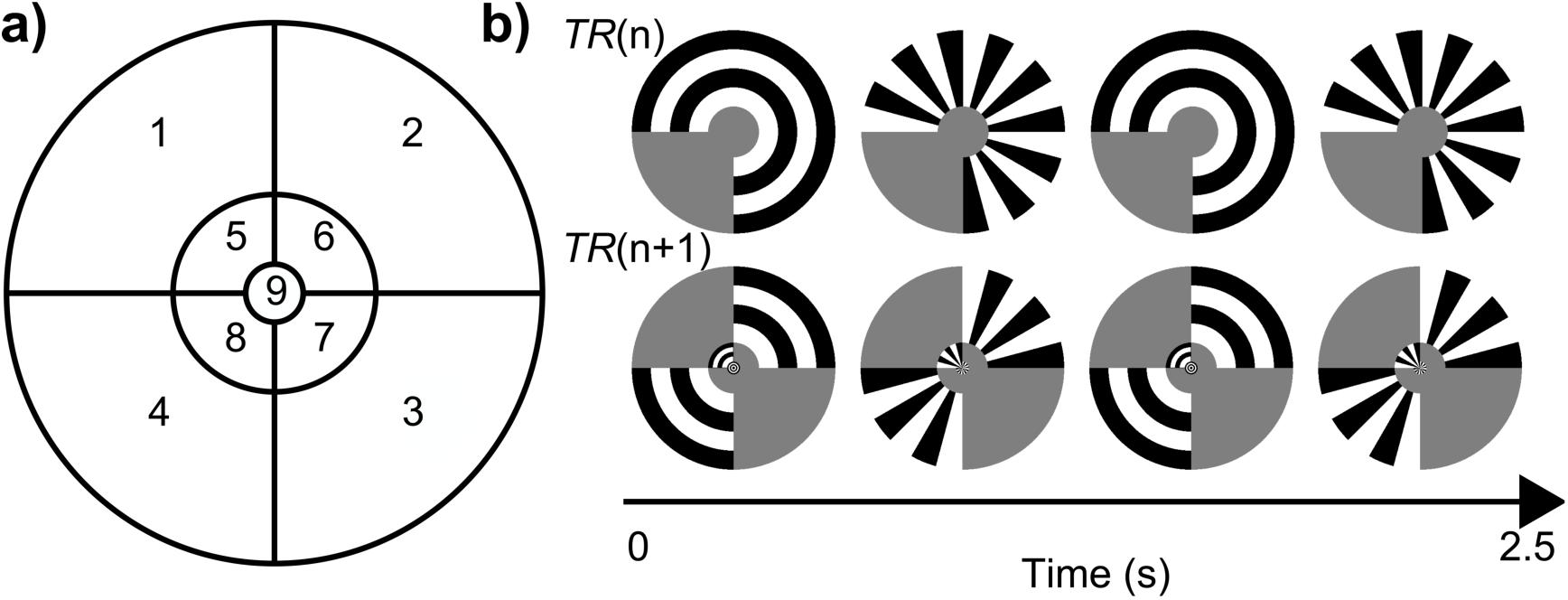
Multifocal stimulation. **(a)** Diagrammatic representation of the nine sector stimulus. These nine patches were presented simultaneously in a pseudo-random m-sequence. **(b)** A representation of the actual stimulus sequence (scaled for V1 magnification). An ‘on’ stimulus comprised the sequence Concentric/Radial/ Concentric/Radial with each image presented for 500 msec. An ‘off’ stimulus was grey (mean luminance). For example, during TR(n) patches 1, 2 and 3 are ‘on’ while the remainder are ‘off’. In the following TR(n + 1) patches 2, 4, 5 and 9 are ‘on’.

Each of the nine patches was treated as a separate stimulus although they were presented simultaneously and in some cases could not be clearly delineated (see Figure 1b). Across time each of these binary stimuli alternated between ‘on’ and ‘off’ states based on the pseudo-random m-sequence. In the ‘on’ state a stimulus patch would produce an alternating pattern across time of concentric-radial-concentric-radial black and white pattern as shown in Figure 1b. This alternation was put in place so as to elicit a strong response from the orientation sensitive visual cortex. Any patch that was in the ‘off’ state had a fixed grey fill (background colour). The beginning of a new repeat time (TR) was used as a trigger for the next iteration of the stimulus sequence.

### M-sequence

Stimuli were presented using Authorware (Macromedia, version 2.2) for Macintosh. The stimulus sequence was determined by an m-sequence iteration of order m, where m = 8. Based on the theory of primitive polynomials modulo 2, the algorithm generates all combinations of 8 bit numbers (except 0) pseudo-randomly, resulting in a sequence of binary strings (where each bit indicates ‘on’ = 1 or ‘off’ = 0 for a given stimulus patch). This pseudo-random method allows maximal decorrelation across patches and balanced presentation across conditions (Press et al., 1986). Vector images were created for all possible combinations of the 8 bit m-sequence. The starting point of the sequence for each region was relatively shifted by the maximum amount possible (256*TR/9), equivalent to a time separation of 70s. As the presentation program (Authorware, Macromedia) iterated through the m-sequence for each patchstimuli with the 9 patches in on or off state, m-sequence the corresponding vector image was displayed.

### Scanning Parameters

FMRI scans were performed at the Brain Research Institute (Melbourne, Australia), using a 3T Signa scanner (General Electric, USA). For each subject a scanning session involved the acquisition of 256 functional volumes (TR = 2.5s; TE = 40ms; NEX = 1; Flip Angle = 60) of T2* EPI images acquired in the axial plane (No. slices = 21; FOV = 240mm; matrix = 128 × 128; slice thickness = 4mm; interslice gap = 1mm). Additionally, high resolution T1 anatomical scans (0.9 mm isovoxel) were taken for coregistration with functional volumes.

### Statistical Analysis

Data was analysed using *BrainVoyager QX* (Brain Innovation, BV, Maastritch, The Netherlands). As a binary m-sequence was used for stimulus presentation, each stimulus patch had an equal amount of ‘on’ and ‘off’ states throughout the entire trial, with this ‘on’/’off’ pattern being decorrelated across patches. This reflects the general rule that more stimuli (in our case patches) require more permutations of the binary string to allow for minimal correlation across stimuli and equality of states within stimuli (Sutter, 1992). Because the stimulus sequence of each of the patches was independent a predicted BOLD signal time course could be built by convolving the sequence with the hemodynamic response function. Thus each of the nine patches (see Fig 1) was analysed separately using a general linear model (derived from the m-sequence) and performing a statistical comparison of ‘on’ *vs*. ‘off’ conditions (corresponding to the first order Wiener kernel). Single subject and groups statistics were performed, with both showing high significance. Statistical thresholding was applied using the false discovery rate (FDR) method as outlined by Genovese et al. (2002), where *q* < 0.005 and *c*(*V*) = 1.

### Simulation

Subsequent to image analysis a post-hoc simulation of the effects of random eye movements during a scanning session was implemented using LabView software (National Instruments, Austin, USA). The effect of “looking” at the wrong segment of the stimulus, given that the segments’ activations is completely decorrelated, is to lower the correlation between stimulus and expected activation. Thus, the whole stimulus image was shifted in vertical and horizontal directions by a Gaussian distribution of random movements (a different position per TR) with a standard deviation of 0.25°. The pixel-by-pixel correlation between the shifted sequence and an unshifted sequence was then calculated.

## Results

A consistent trend in findings was demonstrated across subjects. Only the four large outer peripheral patches showed significant (*p* < .00002 uncorrected, FDR *q* < 0.005) voxel activations (see Figure 2). Significant responses were not obtained from any of the other more central patches. However, as was evident from the conservative FDR level, activation for the four outer patches was highly significant, with no spuriously activated voxels observable in Figure 2. It appears that two sources – likely area V1 and V2 are activated for each stimulus patch.

**Fig. 2.**
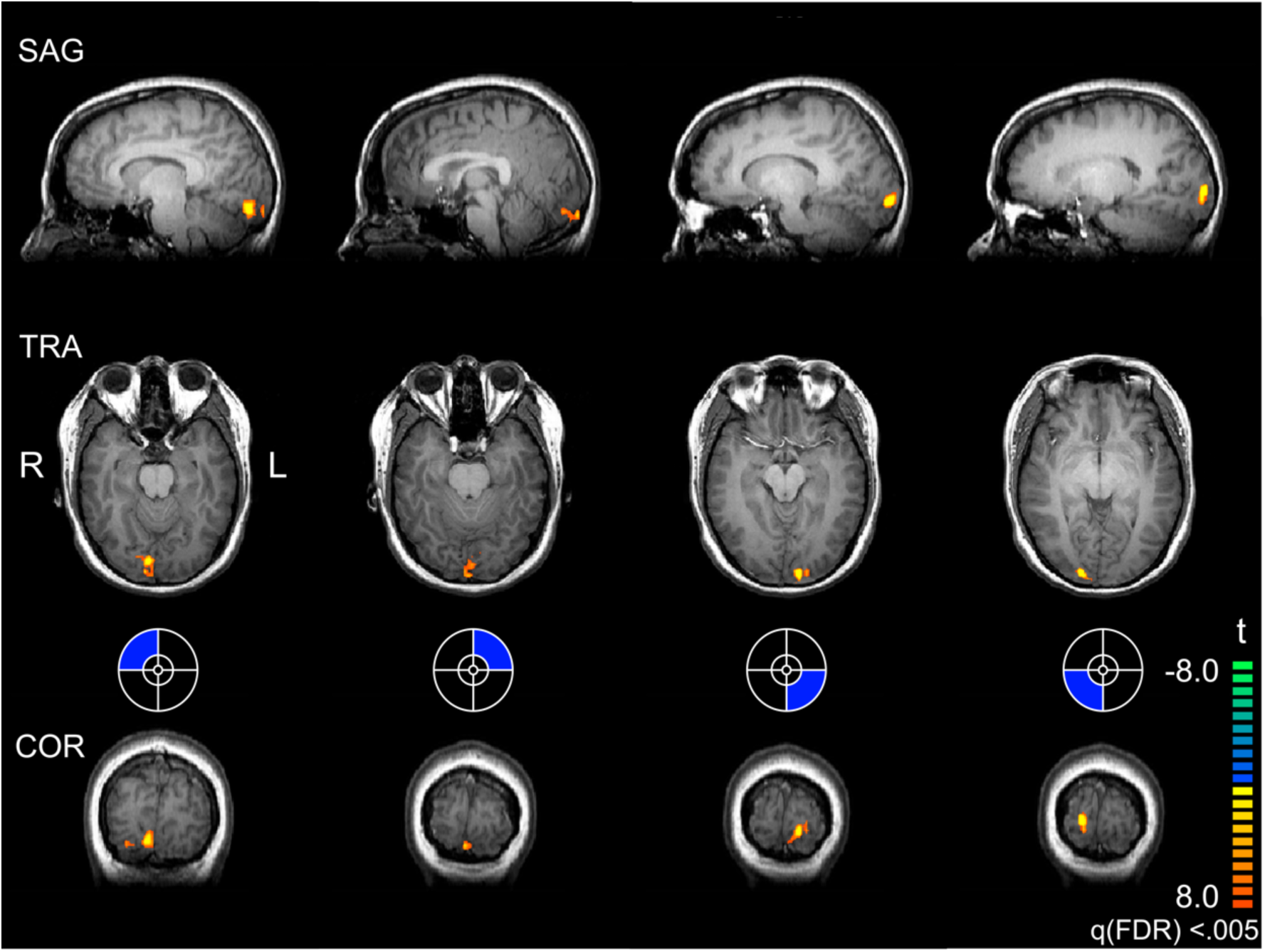
Single subject retinotopy. Example mapping (radiological convention) for a single subject showing excellent retinotopy for the four peripheral quadrants (*p* < 0.00002, uncorrected; q(FDR) < 0.005). (Subject KB). The strength and specificity of the response is evident with no falsely activated voxels observable in the images.

The significant responses in the four peripheral stimulus patches reflect the established retinotopic functional anatomy of V1. It is this retinotopy which provides strong evidence that each patch was measured independently. Our data can therefore be summarised by stating that the outer peripheral patches produced highly significant activations while the remaining inner patches produced non-significant activation. This trend is consistent not only in single subject analysis, but also in group analysis. The averaged event-related response for each of the four peripheral patches across all participants is presented in figure 3. A lag in the BOLD response seen in figure 3 may be due to the rapidity of the stimulus, producing a subsequent delay in response onset or possibly to lateral modulation between stimulated and unstimulated neighbouring regions (Chen et al., 2005). The outer ring patches show very clear activation with signal change ranging from 2.8 – 6.8%.

**Fig. 3.**
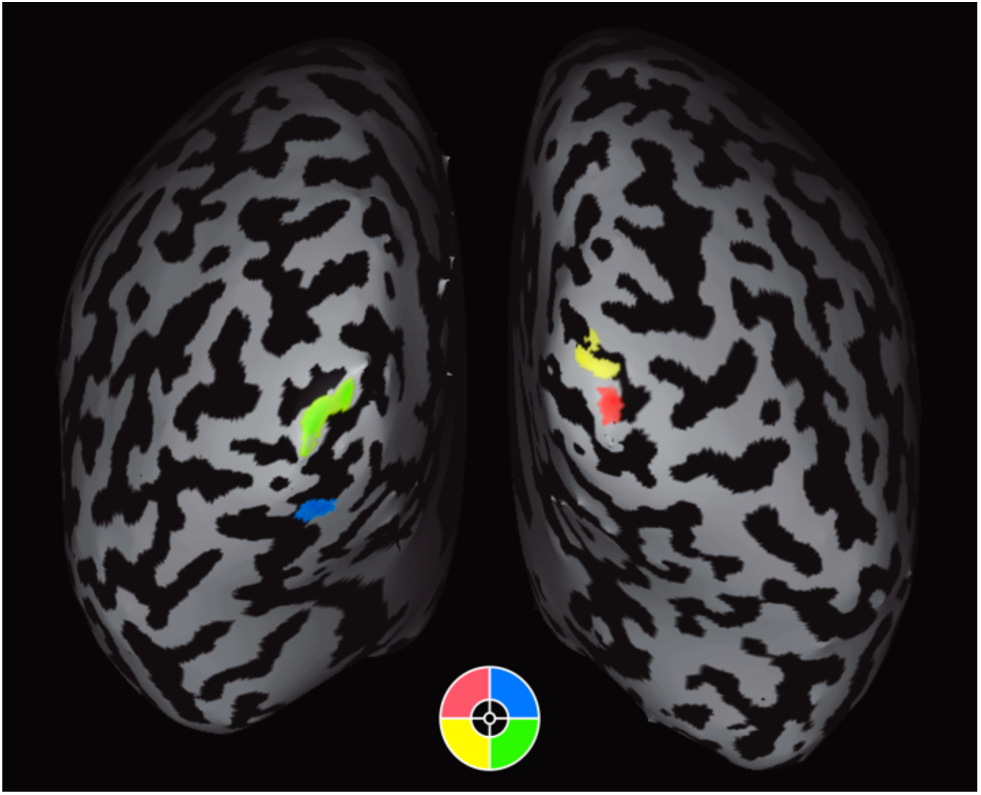
Event related fMRI signal obtained from significantly activated Regions of Interest (ROIs). The curves show the first order kernels (erfMRI_pattern_ – erfMRI_grey_) as means and standard errors for the 6 participants measured from the time of stimulus presentation.

As mentioned earlier, Kellman et al. (2003) used m-sequence based stimulus presentation to investigate second-order nonlinear responses in the visual system. This analysis showed a nonlinear V1 response which was significantly different between foveal and peripheral regions. We also investigated the second-order responses (in the regions providing significant first order response) and found no significant activations across any patches. This is perhaps not surprising on the basis of the slow stimulus rates coupled with a short blank period prior to the start of each stimulus. Indeed, Kellman et al. (2003) report that even a 200ms interstimulus gap greatly diminishes V1 second-order responses.

In light of these results the authors were concerned that while the stimuli were designed to give equal signal to noise on the basis of cortical area of stimulation, eye movements would be more likely to reduce the signal to noise ratio for the regions of smaller area – namely the central regions. Indeed, a number of investigators have already established the fixational sensitivity of both the multifocal ERG (Chisholm et al., 2001;Chu et al., 2006) and multifocal VEP (Menz et al., 2004;Martins et al., 2005). It follows from these findings that multifocal fMRI will be subject to the same sensitivity. Thus, a computational model was set up to test this hypothesis.

### Simulated effects of eye movements

A set of 9 stimulus regions each executing its pseudo-random sequence and being analysed by a perfect observer should give a pixel-by-pixel correlation of 1.0 for each of the regions, if there were no eye movements (see Methods). The effect of fixational inaccuracy was modelled using Gaussian distributed eye position with a standard deviation of 0.25° (reasonable on the basis of human gaze stability against drift and microsaccades (Engbert and Kliegl, 2004)). Additionally, a subsequent study of 3 of the original participants using infra-red eye-movement tracking while observing the stimulus sequence showed an average SD of > 0.25°. The results of the simulation of the effects of eye movements on the effectivity of different parts of the visual field was plotted as a grey scale image of correlation coefficients (see Figure 4).

**Fig. 4.**
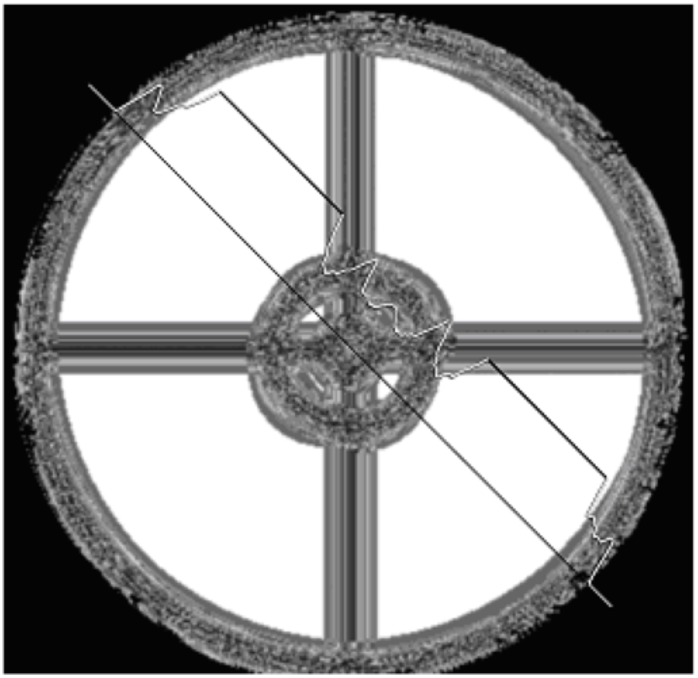
Simulated eyes movements. A correlation map where black = 0 and white = 1. A diagonal section through the image indicates the pixel-by-pixel correlation, showing that the majority of the peripheral patches provide a perfect correlation, suggesting a resilience to eye movements. The same can not be said for the central regions which only indicate small areas of clean signal.

It is clear that the effects of eye movement on the correlation are to reduce the signal derived from the edges of each of the regions. A diagonal section through the image (see Figure 4) demonstrates large areas of perfect correlation in the periphery. The effect on the foveal region was most significant, while small zones of the regions of the next stimulus ring attained better correlations.

## Discussion

The results demonstrate that multifocal fMRI can work in a manner that enables all of the brain volumes scanned to be used for the extraction of signal for different areas, decorrelated by the nature of the stimulus sequence. Even at the rapid rate of stimulation used in this study (2.5 sec versus 7 sec mini-blocks for Vanni et al. (2005)) very strong activations resulted in the periphery. However, the foveal and perifoveal stimulus patches did not produce the expected activations in calcarine striate cortex. Relaxing the threshold for significance did little to alter the number of stimulus areas resulting in significant activation. Also, stimulus size is not the cause as the estimated area of brain activation for the patches of the middle and outer rings were nearly identical. A likely explanation for the eccentricity dependant activations is that eye movements may have caused a dramatic reduction in fMRI signal for stimulus patches subtending small visual angles.

Because the m-sequence relies on a high level of decorrelation across stimuli (patches) and equal state presentations (‘on’/‘off’) within stimuli, eye movements would be expected to affect the outcome. Furthermore, the absence of a stimulus in the foveal segment and the co-occurrence of a stimulus in a neighbouring patch would also provide a strong inducement to move the eyes. The activation resulting from such an inaccurate gaze location would cause the difference between ‘on’ and ‘off’ for the foveal patch analysis to be strongly reduced. Similar arguments apply to the other patches (although the effect on the peripheral edge would tend to be slightly less than that for a boundary separating two active stimulation patches).

The results for the simulation of the effects of eye movements on the multifocal activations recorded provide a convincing account for the lack of activation of the foveal plus the four intermediate peripheral stimulus patches. The magnitude of eye movements employed is within the range of drift and microsaccades shown by normal individuals over the 2.5 s duration of the TR (Engbert and Kliegl, 2003;Horwitz and Albright, 2003). Indeed, it is likely that the nature of the multiple flashed patterns seen by the participants would be an inducement to elicit an eye movement. In the future such visual mapping techniques should take account of both the cortical magnification effects and the effects of eye movements in order to gain the most optimal mapping of visual field regions.

A crucial benefit of multifocal imaging methods is the aspect of efficiency compared with individual scanning of areas. While mentioned by Vanni et al. (2005), the benefit of using all volumes for the analysis of each stimulus patch should be further emphasized. This is especially true if the strength of activation at a particular point in the visual field is required – e.g. if a clinical assessment of potential pathology is being investigated. By comparison, the phase encoded method may not be as efficiently reliable at estimating discrete activation at a particular point in the visual field. The approach of Vanni et al. (2005) to completing the visual field map was through mapping the points of highest activation corresponding to each stimulus patch and smoothing. While this resulted in an excellent mapping, activation level as a function of eccentricity was not discussed. Our research would suggest that eye movements can significantly diminish statistical sensitivity for smaller stimulus patches despite equivalent cortical areas of activation.

## Conclusion

The standard multifocal paradigm of electrophysiology can be effectively applied to fMRI, especially if seeking a method for efficiently establishing the functional integrity at any point on the visual map. However, when many small visual field patches are to be used as visual stimuli, the results are very dependent on proper ocular fixation. Such control may be achieved in the future with retinally stabilized stimulus presentation.

## Acknowledgments

Part of the data reported here was previously presented at Human Brain Mapping (2004), in Budapest, Hungary. The research was supported by a Discovery Project DP0345767 from the Australian Research Council.

## References

Baker, S., Baseler, H., Klein, S., and Carney, T. (2006). Localizing sites of activation in primary visual cortex using visual-evoked potentials and functional magnetic resonance imaging. J Clin Neurophysiol 23, 404–415.

Buracas, G.T., and Boynton, G.M. (2002). Efficient design of event-related fMRI experiments using M-sequences. Neuroimage 16, 801–813.

Chen, C.C., Tyler, C.W., Liu, C.L., and Wang, Y.H. (2005). Lateral modulation of BOLD activation in unstimulated regions of the human visual cortex. Neuroimage 24, 802–809.

Chisholm, J.A., Keating, D., Parks, S., and Evans, A.L. (2001). The impact of fixation on the multifocal electroretinogram. Doc Ophthalmol 102, 131–139.

Chu, P.H., Chan, H.H., and Leat, S.J. (2006). Effects of unsteady fixation on multifocal electroretinogram (mfERG). Graefes Arch Clin Exp Ophthalmol 244, 1273–1282.

Crewther, S.G., Crewther, D.P., Klistorner, A., and Kiely, P.M. (1999). Development of the magnocellular VEP in children: implications for reading disability. Electroencephalogr Clin Neurophysiol Suppl 49, 123–128.

De Zwart, J.A., Silva, A.C., Van Gelderen, P., Kellman, P., Fukunaga, M., Chu, R., Koretsky, A.P., Frank, J.A., and Duyn, J.H. (2005). Temporal dynamics of the BOLD fMRI impulse response. Neuroimage 24, 667–677.

Deyoe, E.A., Carman, G.J., Bandettini, P., Glickman, S., Wieser, J., Cox, R., Miller, D., and Neitz, J. (1996). Mapping striate and extrastriate visual areas in human cerebral cortex. Proc Natl Acad Sci U S A 93, 2382–2386.

Duncan, R.O., and Boynton, G.M. (2003). Cortical magnification within human primary visual cortex correlates with acuity thresholds. Neuron 38, 659–671.

Engbert, R., and Kliegl, R. (2003). Microsaccades uncover the orientation of covert attention. Vision Res 43, 1035–1045.

Engbert, R., and Kliegl, R. (2004). Microsaccades keep the eyes’ balance during fixation. Psychol Sci 15, 431–436.

Engel, S.A., Glover, G.H., and Wandell, B.A. (1997). Retinotopic organization in human visual cortex and the spatial precision of functional MRI. Cereb Cortex 7, 181–192.

Genovese, C.R., Lazar, N.A., and Nichols, T. (2002). Thresholding of statistical maps in functional neuroimaging using the false discovery rate. Neuroimage 15, 870–878.

Hansen, K.A., David, S.V., and Gallant, J.L. (2004). Parametric reverse correlation reveals spatial linearity of retinotopic human V1 BOLD response. Neuroimage 23, 233–241.

Henriksson, L., Karvonen, J., Salminen-Vaparanta, N., Railo, H., and Vanni, S. (2012). Retinotopic maps, spatial tuning, and locations of human visual areas in surface coordinates characterized with multifocal and blocked FMRI designs. PLoS One 7, e36859.

Hood, D.C. (2000). Assessing retinal function with the multifocal technique. Prog Retin Eye Res 19, 607–646.

Horwitz, G.D., and Albright, T.D. (2003). Short-latency fixational saccades induced by luminance increments. J Neurophysiol 90, 1333–1339.

James, A.C. (2003). The pattern-pulse multifocal visual evoked potential. Invest Ophthalmol Vis Sci 44, 879–890.

Kellman, P., Gelderen, P., De Zwart, J.A., and Duyn, J.H. (2003). Method for functional MRI mapping of nonlinear response. Neuroimage 19, 190–199.

Klistorner, A., Crewther, D.P., and Crewther, S.G. (1997). Separate magnocellular and parvocellular contributions from temporal analysis of the multifocal VEP. Vision Res 37, 2161–2169.

Liu, T.T. (2004). Efficiency, power, and entropy in event-related fMRI with multiple trial types. Part II: design of experiments. Neuroimage 21, 401–413.

Liu, T.T., Frank, L.R., Wong, E.C., and Buxton, R.B. (2001). Detection power, estimation efficiency, and predictability in event-related fMRI. Neuroimage 13, 759–773.

Maddess, T., Bedford, S., James, A.C., and Rose, K.A. (1997). A multiple-frequency, multiple-region pattern electroretinogram investigation of non-linear retinal signals. Aust N Z J Ophthalmol 25 Suppl 1, S94–97.

Martins, A., Klistorner, A., Graham, S., and Billson, F. (2005). Effect of fixation tasks on multifocal visual evoked potentials. Clin Experiment Ophthalmol 33, 499–504.

Menz, M., Sutter, E., and Menz, M. (2004). The effect of fixation instability on the multifocal VEP. Doc Ophthalmol 109, 147–156.

Nishiyama, T., Ohde, H., Haruta, Y., Mashima, Y., and Oguchi, Y. (2004). Multifocal magnetoencephalogram applied to objective visual field analysis. Jpn J Ophthalmol 48, 115–122.

Press, W., Flannery, B., Teukolsky, S., and Vetterling, W. (1986). Numerical Recipes in C: The Art of Scientific Computing.

Schneider, W., Noll, D.C., and Cohen, J.D. (1993). Functional topographic mapping of the cortical ribbon in human vision with conventional MRI scanners. Nature 365, 150–153.

Sereno, M.I., Dale, A.M., Reppas, J.B., Kwong, K.K., Belliveau, J.W., Brady, T.J., Rosen, B.R., and Tootell, R.B. (1995). Borders of multiple visual areas in humans revealed by functional magnetic resonance imaging. Science 268, 889–893.

Sutter, E.E. (1992). “A deterministic approach to nonlinear systems analysis.,” in Nonlinear Vision, eds. R.B. Pinter & B. Nabet. (Cleveland: CRC Press), 171–220.

Tabuchi, H., Yokoyama, T., Shimogawara, M., Shiraki, K., Nagasaka, E., and Miki, T. (2002). Study of the visual evoked magnetic field with the m-sequence technique. Invest Ophthalmol Vis Sci 43, 2045–2054.

Vanni, S., Henriksson, L., and James, A.C. (2005). Multifocal fMRI mapping of visual cortical areas. Neuroimage 27, 95–105.

Vanni, S., Warnking, J., Dojat, M., Delon-Martin, C., Bullier, J., and Segebarth, C. (2004). Sequence of pattern onset responses in the human visual areas: an fMRI constrained VEP source analysis. Neuroimage 21, 801–817.

